# Pro-Cognitive Effects of IgM Isotype Anti-NMDAR1 Autoantibodies in Mice

**DOI:** 10.64898/2026.04.24.720689

**Authors:** Jenna M. DeWit, Tina Tebyanian, Alexandra Unapanta, Melonie N. Vaughn, Susan B. Powell, Victoria B. Risbrough, Xianjin Zhou

## Abstract

Natural anti-NMDAR1 autoantibodies are present at varying levels in the general human population, but their effects on cognitive function remain unclear. Recent human studies reported significant associations between higher blood levels of natural anti-NMDAR1 autoantibodies and potential neuroprotective outcomes in Alzheimer’s disease, traumatic brain injury–associated depression and PTSD symptoms, and schizophrenia. However, whether these natural autoantibodies play a causal role in emotional and cognitive function has not been investigated. Since natural autoantibodies in human blood are predominantly of the IgM isotype, we immunized Aicda mutant mice to produce only IgM isotype anti-NMDAR1 autoantibodies without IgG and IgA isotypes. Mice were tested for sensorimotor gating and conditioned fear and extinction, cross species measures of information processing and emotional memory, respectively. Mice with higher levels of IgM anti-NMDAR1 autoantibodies exhibited significantly increased sensorimotor gating and improved fear extinction recall compared with mice with baseline levels of these autoantibodies. These findings indicate that IgM anti-NMDAR1 autoantibodies are pro-cognitive, unlike previous reports of poor cognition associated with IgG anti-NMDAR1 autoantibodies. Together, these studies suggest that IgM may hold therapeutic potential for a range of neurodegenerative, neurological, and psychiatric disorders.

## Introduction

High titers of IgG anti-NMDAR1 autoantibodies in brain are involved in the anti-NMDAR1 encephalitis that exhibits psychosis, impaired memory, and many other psychiatric symptoms in addition to neurological symptoms (Dalmau, 2016). On the other hand, low titers of natural anti-NMDAR1 autoantibodies have been detected in the blood of ∼10% of the general human population using cell-based assays (Castillo-Gomez *et al*, 2017; Hammer *et al*, 2014; Jezequel *et al*, 2017; Pan *et al*, 2019). Their effects on human cognitive function are unclear (Zhou, 2021, 2023). To investigate potential effects of blood circulating IgG anti-NMDAR1 autoantibodies on behavioral phenotypes, we generated mice carrying low titers of anti-NMDAR1 autoantibodies, predominantly IgG isotype, in blood against a single P2 antigenic epitope of mouse NMDAR1 using active immunization (Yue *et al*, 2021). Mice carrying the IgG anti-NMDAR1 P2 autoantibodies are healthy and display no differences in locomotion, sensorimotor gating, memory, and depression-like phenotypes compared to controls. Presence of blood IgG anti-NMDAR1 autoantibodies, however, is sufficient to significantly impair spatial working memory with intact blood-brain barriers (BBB) (Yue *et al*., 2021).

To more accurately quantify blood anti-NMDAR1 autoantibody levels, we recently developed a novel cross-species immunoassay (Vaughn *et al*, 2025). Using this immunoassay, we found that higher levels of blood natural anti-NMDAR1 autoantibodies are significantly associated with slower cognitive decline in patients with Alzheimer’s disease (Zhou, 2026), as well as less severe depression and post-traumatic stress disorder symptoms in Marines with traumatic brain injuries (Vaughn *et al*, 2026). Consistent with our findings, schizophrenia patients who are seropositive for anti-NMDAR1 autoantibodies exhibit less severe negative symptoms and better social functioning than seronegative patients (Hansen *et al*, 2024; Luykx *et al*, 2024). In humans, natural autoantibodies are mostly IgM isotype with a minor portion of IgA and IgG3 (Elkon & Casali, 2008; Lobo, 2016). Thus, we are interested in investigating whether IgM isotype ant-NMDAR1 autoantibodies may be beneficial for cognitive function.

Activation-induced cytidine deaminase (*Aicda*) is essential for producing somatic hypermutation (SHM) of Ig genes and class switch recombination (CSR) in generation of IgG and IgA antibodies. Knockout of the *Aicda* gene prevents production of IgG and IgA antibodies, but not IgM antibodies (Nilaratanakul *et al*, 2018). Here, we report the successful generation of IgM isotype anti-NMDAR1 autoantibodies against NMDAR1 P2 antigen in Aicda homozygous mice in the absence of IgG and IgA isotypes. We found that Aicda mice producing IgM anti-NMDAR1 P2 autoantibodies are healthy and exhibit enhanced sensorimotor gating and cognitive performance, supporting their beneficial effects suggested by human association studies.

## Materials and Methods

### Mouse strain

*Aicda* homozygous mice (B6.129P2-*Aicda*^*tm1(cre)Mnz*^/J; JAX), backcrossed to C57 BL/6 background for 11 generations, were purchased from Jackson Labs (Bar Harbor, ME) at 8-week-old and housed in a climate-controlled animal colony with a reversed day/night cycle. Food (Harlan Teklab, Madison, WI) and water were available *ad libitum*, except during behavioral testing. Active immunization was conducted after a week of acclimation to the facility. Blood was taken from mouse tail vein one month after immunization to detect production of IgM anti-NMDAR1 autoantibodies. Behavioral testing began one week after blood collection. All testing procedures were approved by UCSD Animal Care and Use Committee (S09179) prior to the onset of the experiments. Mice were maintained in American Association for Accreditation of Laboratory Animal Care approved animal facilities at UCSD campus, which meet all Federal and State requirements for animal care.

### Active immunization

Active immunization of Aicda homozygous mutant mice were conducted as described in our previous publication (Yue *et al*., 2021). Peptide P2 (TIHQEPFVYVKPTMSDGTCK), located in the ligand binding domain of mouse NMDAR1, was synthesized by *Biomatik. Mycobacterium tuberculosis*, H37 RA (Difco) and Incomplete Freund’s Adjuvant (Bacto) were purchased for active immunization.

#### Mouse Cohort 1

Complete Freund’s Adjuvant (CFA) was used for immunization of mouse cohort 1. P2 peptide was dissolved in PBS at a concentration of 4 mg/ml. An equal volume of the P2 solution is completely mixed with complete Freund’s adjuvant (containing 4 mg/ml desiccated *M. Tuberculosis*, H37 RA in Incomplete Freund’s Adjuvant) to generate a thick emulsion. Twenty mice were injected with 100 ul of emulsion subcutaneously divided equally at 2 sites on mouse flank. For the injection of 20 control mice, the same emulsion was generated but without the P2 peptide.

#### Mouse Cohort 2

Incomplete Freund’s Adjuvant was used for immunization of mouse cohort 2. No Mycobacterium tuberculosis H37RA was included in the emulsion. A total of 20 mice received emulsion containing the P2 antigen, and 20 control mice received emulsion without the P2 antigen.

#### Mouse Cohort 3

Complete Freund’s Adjuvant was used for immunization of mouse cohort 3; however, the CFA formulation contained only 50% of the standard amount of Mycobacterium tuberculosis H37RA (2 mg/mL desiccated M. tuberculosis H37RA in Incomplete Freund’s Adjuvant). A total of 33 mice received emulsion containing the P2 antigen, and 29 control mice received emulsion without the P2 antigen.

### Quantification of IgM and IgG anti-NMDAR1 autoantibodies

IgM and IgG isotypes of anti-NMDAR1 autoantibodies were quantified using our recently developed immunoassay (Vaughn *et al*., 2025) with some modifications. Protein AG (Prospec, cat. PRO-646-B) was incubated for 1 hour at room temperature with either Rabbit anti-mouse IgM (Jackson ImmunoResearch, 315-005-020) or Rabbit anti-mouse IgG (Jackson ImmunoResearch, 315-005-008). After this incubation, 4 μl of rabbit serum (Jackson ImmunoResearch, 011-000-120) was added to saturate all remaining Protein AG binding sites. In parallel, the NMDAR1-GLUC probe was incubated with 2 μl of mouse serum, along with diluted goat serum used as the background for the negative control. Seven microliters of the Protein-AG/antibody suspension were added to 5 μl of the mouse serum/NMDAR-GLUC mixture and incubated for 1.5 hours at room temperature. Positive mouse serum samples with different levels of IgM or IgG anti-NMDAR1 autoantibodies were used as positive controls and references for normalization between different 96-well plates (Greiner 96-well Flat Bottom Black Polystyrene plate, Cat. No.: 655097). Each mouse serum sample was supplemented with the same amount of goat serum as in the negative control for the immunoassay. *Gaussia* luciferase substrate (ThermoFisher, cat. 16160; Pierce™ *Gaussia* Luciferase Glow Assay Kit) was used for quantification on Tecan Spark plate reader. After subtracting the Relative Light Units (RLU) of the negative control, the remaining RLU will be the levels of anti-NMDAR1 autoantibodies in individual samples.

### Prepulse inhibition

Mouse startle reactivity and prepulse inhibition (PPI) were measured with startle chambers (SR-LAB, San Diego Instruments, San Diego, CA) as described in our previous publications (Ji *et al*, 2013).

### Fear conditioning

Fear conditioning procedures followed previously published protocols (Gresack *et al*, 2010; Toth *et al*, 2014; Yue *et al*., 2021) and were conducted in four automated conditioning chambers (Med Associates Inc., St. Albans, VT). Freezing behavior was quantified using Video Freeze software (Med Associates Inc.) with a motion index threshold of 30 frames and a minimum freeze duration of 18 frames (Anagnostaras *et al*, 2010). Foot shocks were delivered through 36 stainless steel floor rods.

#### Day 1 – Acquisition

After a 60 min habituation period in an adjacent room, mice were placed into the conditioning chamber. Following a 2 min acclimation period, mice received five tone shock pairings consisting of a 20 sec tone (CS; 75 dB, 4 kHz) co-terminating with a 1 sec, 0.7 mA foot shock (US). Inter-trial intervals were 40 sec.

Freezing was scored during each tone and during the 40 sec post-shock interval. Mice were returned to their home cages 40 sec after the final shock. Chambers were cleaned with 70% ethanol between sessions.

#### Day 2 – Context Memory

Mice were re-exposed to the original conditioning chamber under identical contextual conditions (e.g. shock grids, 70% EtOH, lights on). After placement in the chamber, freezing was recorded for 16 min in the absence of tones or shocks.

#### Day 3 – Cued Fear and Extinction

To minimize contextual cues, chamber context was altered across tactile (solid Plexiglas floor), olfactory (1% almond extract in 30% EtOH for cleaning), and visual (lights off) dimensions. After a 2-min acclimation period without tones (“pre-tone”), mice were presented with 32 CS tones to assess cued fear and extinction learning.

#### Day 4 – Extinction Recall

Twenty-four hours after extinction training, mice were returned to the altered context. Following a 2-min acclimation period without tones, 12 CS tones were presented to evaluate recall of extinction learning.

### Statistical analysis

R programming was used for statistical analyses. Analysis of variance (ANOVA) was used for analysis of IgM anti-NMDAR1 autoantibody levels with antigen P2, sex, and cohort as between-subject factors. ANOVA with anti-NMDAR1 autoantibody and sex as between-subjects factors, block and prepulse intensity as within-subjects factors were performed on the %PPI data. For fear conditioning, ANOVA was conducted with anti-NMDAR1 autoantibody and sex as between-subjects factors and CS or time block as a within-subject factor on freezing percentages. When there is no sex interaction, both sexes were combined for analysis. *Post hoc* analyses were carried out using Tukey’s test. Alpha level was set to 0.05.

## Results

To generate mice carrying IgM anti-NMDAR1 autoantibodies, we immunized Aicda-deficient mice with the NMDAR1 P2 antigen that has been shown to robustly induce anti-NMDAR1 autoantibody production in wildtype C57BL/6 mice (Yue *et al*., 2021). Three independent large cohorts of Aicda mutant mice (Table I) were used to evaluate the efficiency of IgM anti-NMDAR1 autoantibody induction using the P2 antigen formulated with either CFA containing 4 mg/ml desiccated M. tuberculosis H37RA, Incomplete Freund’s Adjuvant, or a reduced-strength CFA containing 2 mg/ml desiccated M. tuberculosis H37RA. IgM anti-NMDAR1 autoantibodies were successfully generated with all three P2 antigen formulations (Figure 1). As expected, no IgG anti-NMDAR1 autoantibodies were detected in Aicda mutant mice, regardless of whether they were immunized with the P2 antigen or not (Figure 1A). Significant effects were observed for the P2 antigen (F(1,127) = 94.695, p < 2 × 10^−16^), cohort (F(2,127) = 5.01, p = 0.0081), and the interaction between P2 and cohort (F(2,127) = 4.40, p = 0.0142) (Figure 1D). IgM anti-NMDAR1 autoantibodies appeared to be most effectively induced when the P2 antigen was formulated with the reduced-strength CFA containing 2 mg/ml desiccated M. tuberculosis H37RA.

**Table 1.** Three independent cohorts of Aicda mutant mice.

| **Cohort** | **Control Immunization** |  | **P2 Immunization** |  | **Total Mice** |
| --- | --- | --- | --- | --- | --- |
|  | Female | Male | Female | Male |  |
| 1 | 10 | 10 | 10 | 10 | 40 |
| 2 | 10 | 10 | 10 | 10 | 40 |
| 3 | 13 | 16 | 17 | 16 | 62 |

**Figure 1.**
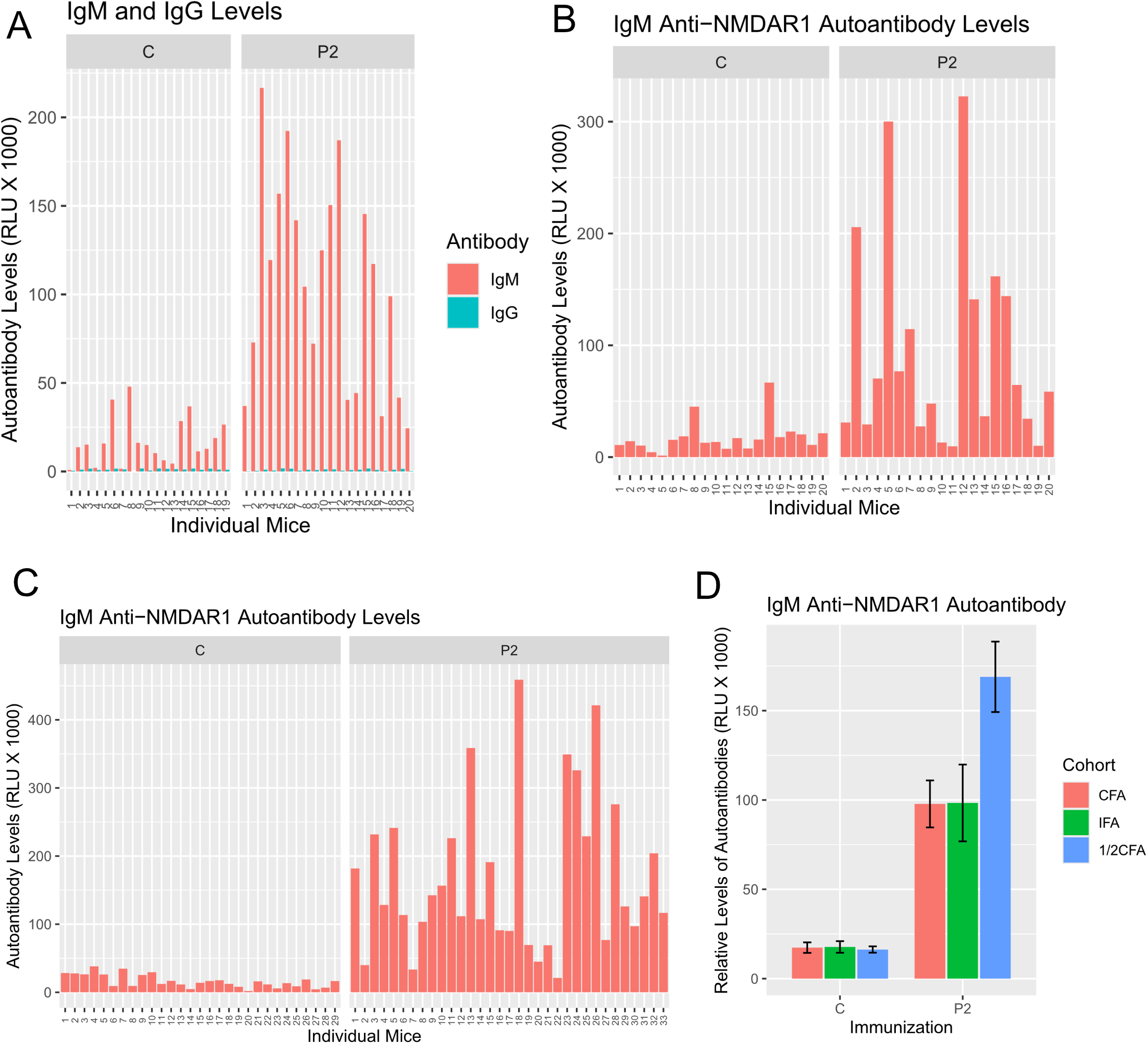
Generation of blood IgM anti-NMDAR1 autoantibodies against NMDAR1 P2 antigen in Aicda mutant mice by active immunization. Mice were immunized with either Complete Freund’s Adjuvant (CFA) or Incomplete Freund’s Adjuvant in the absence (Control) or presence (P2) of the NMDAR1 P2 antigen. Quantification of blood anti-NMDAR1 autoantibodies were conducted one month after immunization. **(A)** Mouse cohort 1 was immunized with CFA (containing 4 mg/ml desiccated M. tuberculosis, H37RA) with or without the P2 peptide. IgM anti-NMDAR1 autoantibodies were induced in mice immunized with CFA plus P2, whereas only baseline levels of IgM autoantibodies were detected in control mice receiving CFA alone. No IgG anti-NMDAR1 autoantibodies were detected in either group regardless of P2 immunization. **(B)** Mouse cohort 2 was immunized with Incomplete Freund’s Adjuvant with or without the P2 peptide. IgM anti-NMDAR1 autoantibodies were induced in mice immunized with P2, whereas only baseline levels of IgM autoantibodies were detected in control mice. **(C)** Mouse cohort 3 was immunized with CFA (containing 2 mg/ml desiccated M. tuberculosis, H37RA) with or without the P2 peptide. IgM anti-NMDAR1 autoantibodies were induced in mice immunized with CFA plus P2, whereas only baseline levels of IgM autoantibodies were detected in control mice receiving CFA alone. **(D)** A significant interaction between P2 and cohort was detected (F(2,127) = 4.40, p = 0.0142). Levels of IgM anti-NMDAR1 autoantibodies in cohort 3 were significantly higher than in either cohort 1 or cohort 2, suggesting that CFA containing 2 mg/ml desiccated M. tuberculosis H37RA is more effective at inducing IgM responses. CFA: Complete Freund’s Adjuvant (containing 4 mg/ml desiccated M. tuberculosis, H37RA); IFA: Incomplete Freund’s Adjuvant, 1/2CFA: Complete Freund’s Adjuvant (containing 2 mg/ml desiccated M. tuberculosis, H37RA). Data were presented as Mean+SEM.

All immunized Aicda mice are healthy and display no differences in locomotor activity between mice immunized with or without the NMDAR1 P2 antigen (data not shown). Because NMDA receptor signaling modulates sensorimotor gating, we assessed prepulse inhibition (PPI) to compare sensorimotor gating between mice immunized with or without the P2 antigen. A trend toward increased PPI was observed in cohorts 1 and 2, respectively. A significant increase in PPI was observed in mice immunized with the P2 antigen compared with control mice (F(1,74) = 4.932, p = 0.029) when data from both cohorts were combined (Supplemental Figure 1A). There were no differences in startle (F(1,67)=0.044, p=0.835) or startle habituation (F(3, 201)=0.787, p=0.502) between the two groups (Supplemental Figure 1B). Increased PPI was replicated in a larger cohort 3. PPI was significantly increased in mice immunized with the P2 antigen compared with control mice (F(1, 58)=4.978, p=0.0295; Supplemental Figure 1C). Surprisingly, startle responses were significantly suppressed in mice immunized with the P2 antigen compared with control mice (F(1,58) = 12.328, p = 0.0009; Supplemental Figure 1D). No difference in startle habituation was observed between the two groups. When all three cohorts were combined, we found a significant increase in PPI in mice immunized with the P2 antigen compared with controls (F(1,128) = 9.527, p = 0.0025; Figure 2A). *Post hoc* analysis revealed significantly increased PPI in P2-immunized mice across all three prepulse intensities. A significant suppression of startle responses was also observed in P2-immunized mice (F(1,128) = 5.13, p = 0.025), whereas startle habituation did not differ between the groups (Figure 2B).

**Figure 2.**
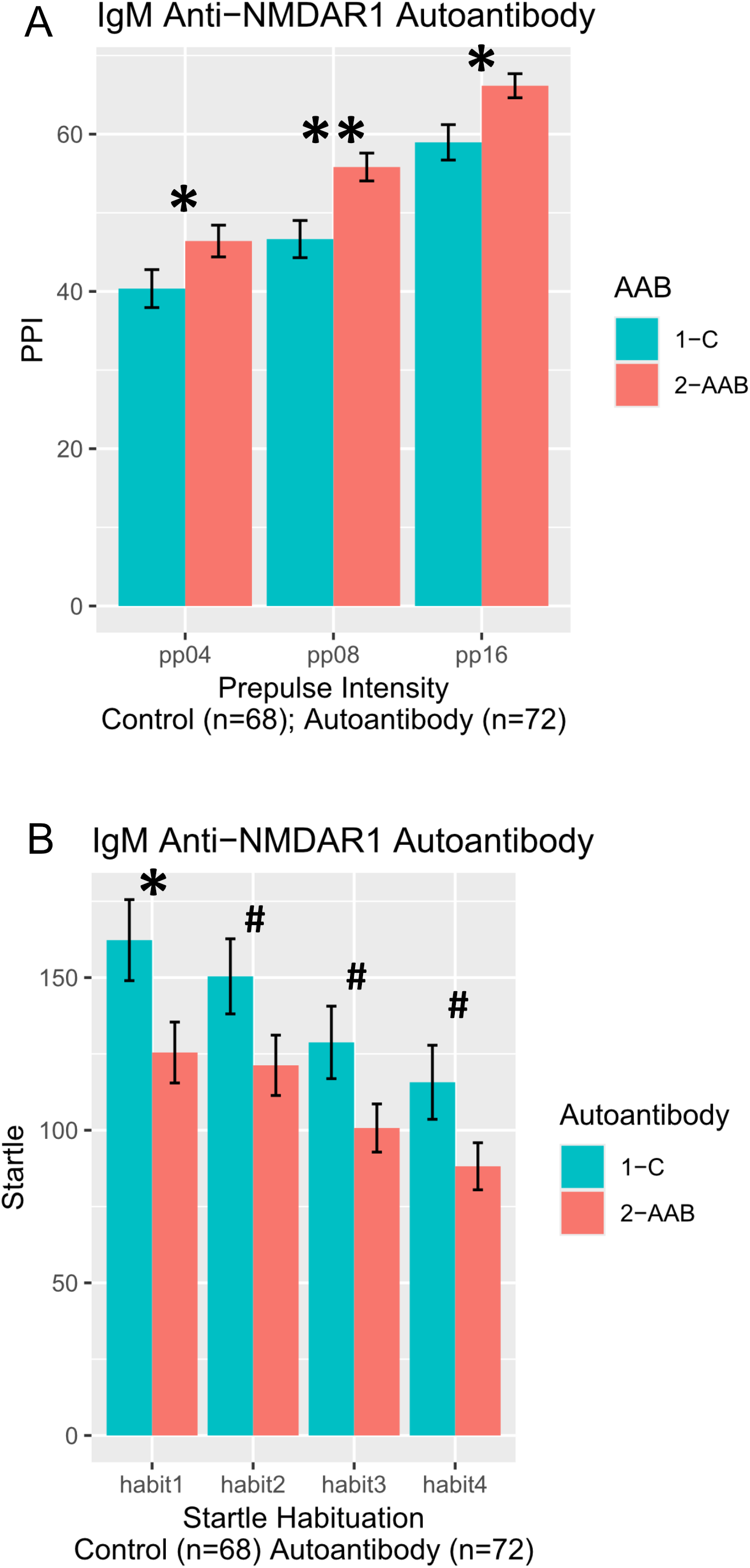
Enhancement of prepulse inhibition in Aicda mutant mice immunized with NMDAR1 P2 antigen. All 3 cohorts of mice were combined for analysis. **(A)** Mice immunized with the P2 antigen showed a significant increase in PPI compared with controls (F(1,128) = 9.527, p = 0.0025). Post hoc analyses revealed that PPI was significantly elevated in P2-immunized mice across all three prepulse intensities. **(B)** P2 immunization significantly suppressed startle responses (F(1,128) = 5.13, p = 0.025), but startle habituation did not differ between two groups. C: Control; AAB: IgM anti-NMDAR1 autoantibodies. Data were presented as Mean+SEM. p value: ** < 0.01; * < 0.05;^#^ < 0.1.

Mice were further tested using fear conditioning to assess fear learning, memory, and extinction. Because autoantibody effect sizes were small, all three mouse cohorts were combined for analysis. No differences were observed between the two groups in Day 1 fear acquisition during the tone (CS; Figure 3A) or during the post-shock period (Figure 3B). On Day 2, mice were returned to the conditioning chamber for a 16-minute context re-exposure session (Figure 3C); freezing during the first 4 minutes was used for calculation of contextual memory (Figure 3D). No differences were observed between the two groups. On Day 3, cued memory and fear extinction were evaluated (Figure 4A). A significant fear extinction across CS sessions was observed (F(31, 4185) = 60.696, p < 2 × 10^−16^). Autoantibody X CS interaction was not significant (F(31, 4185) = 1.238, p = 0.17). Freezing percentages from CS1 were used to quantify cued memory (Figure 4B). No cued memory difference was observed between the two groups. On Day 4, fear extinction recall was examined (Figure 4C). A significant fear extinction across CS sessions was observed (F(11, 1485) = 60.99, p < 2 × 10^−16^). No significant interaction was observed between autoantibody and CS sessions. Freezing percentages from CS1 were used to quantify fear extinction recall (Figure 4D). A significant better fear extinction recall was observed in the P2-immunized mice than control mice (F(1, 137)=5.09, p=0.0256).

**Figure 3.**
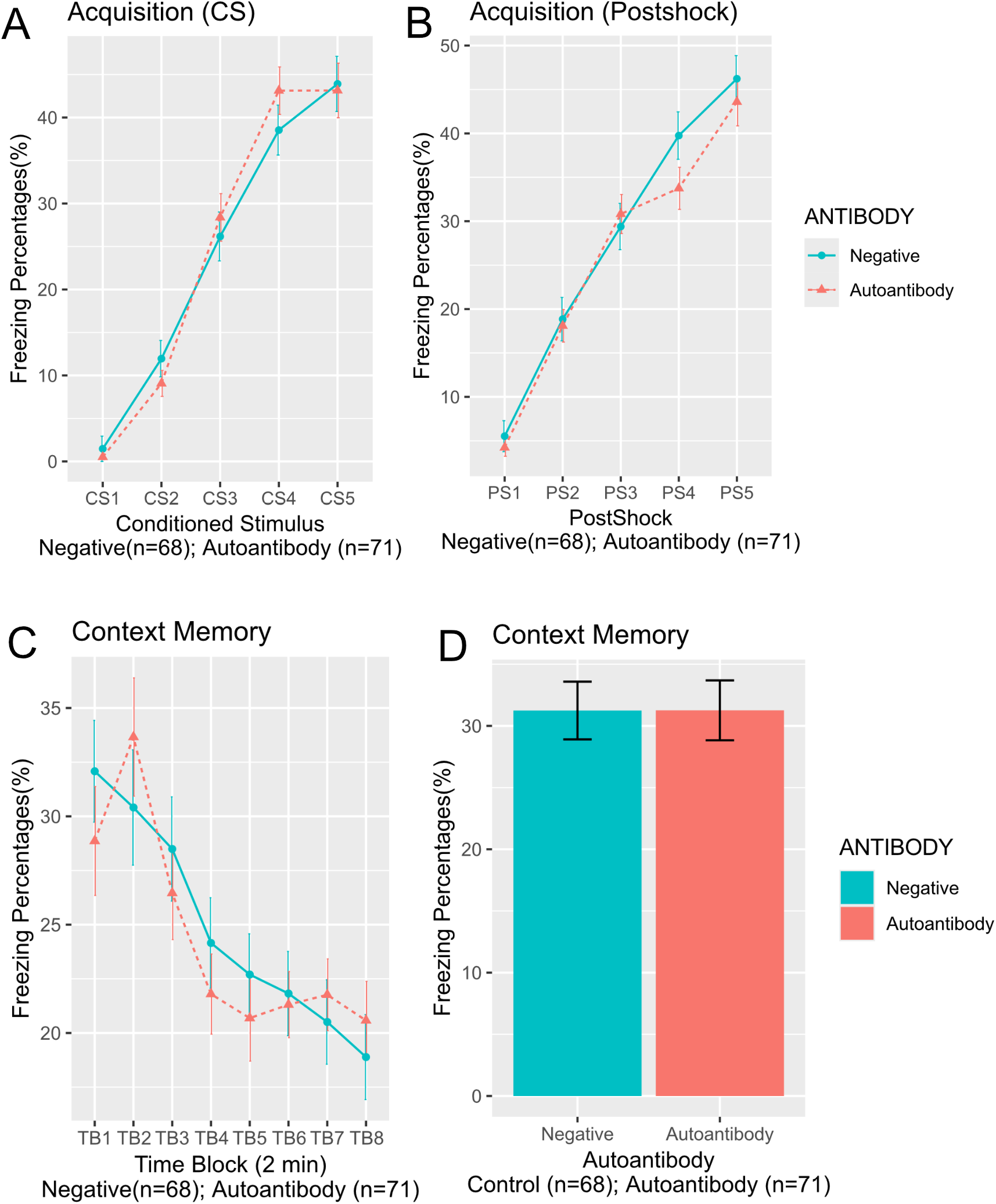
No differences in fear acquisition and contextual memory. All 3 cohorts of mice were combined for analysis. No differences were observed between the two groups in Day 1 fear acquisition during the tone (**A**) or during the post-shock period **(B). (C)** On Day 2, mice underwent a 16-minute context re-exposure session to assess contextual fear memory. **(D)** Contextual memory was assessed by measuring freezing behavior during the first four minutes, and no differences were observed between the two groups. Negative: Control mice; Autoantibody: IgM anti-NMDAR1 autoantibody. Data were presented as Mean+SEM.

**Figure 4.**
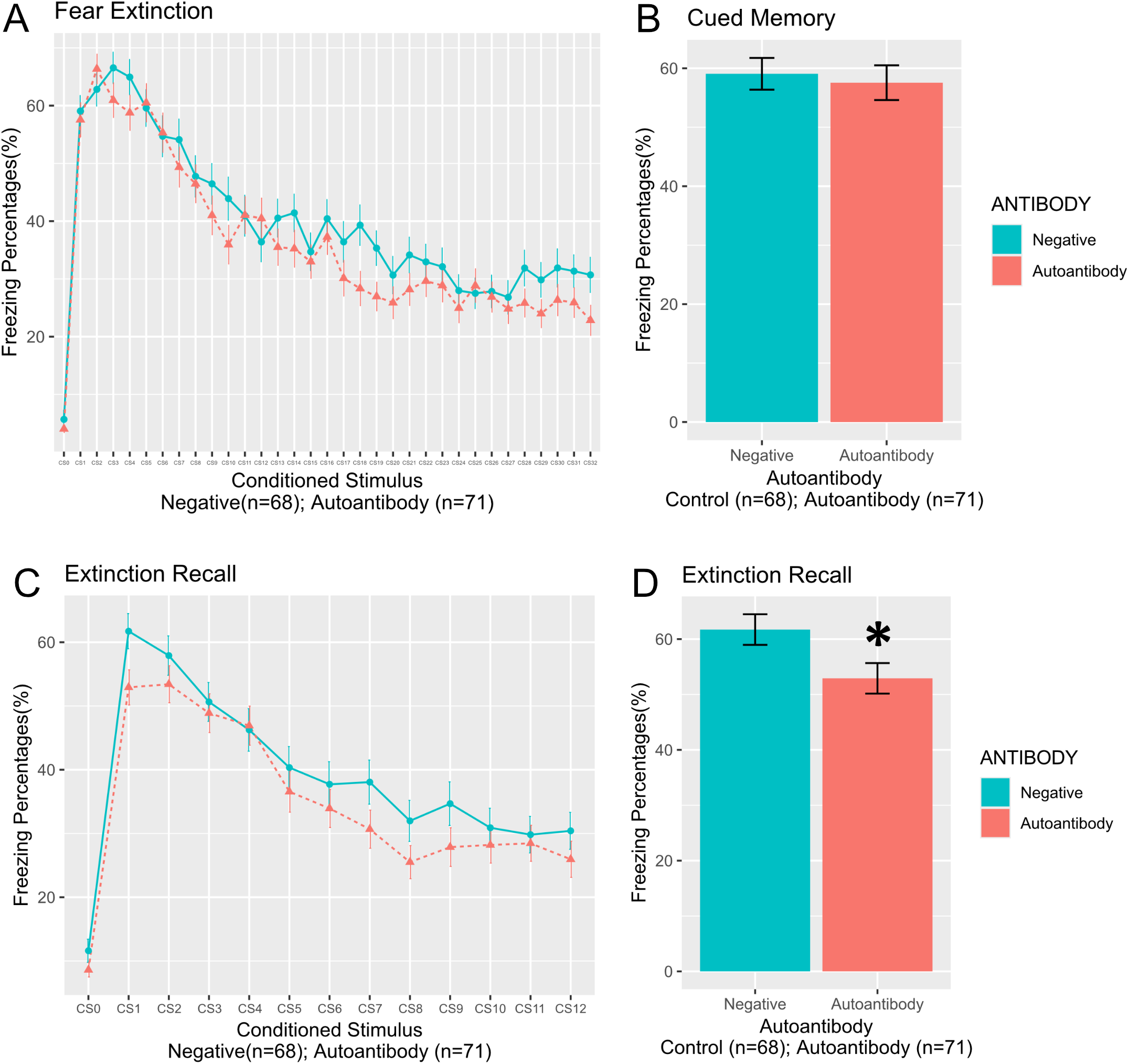
Enhanced fear extinction recall in Aicda mutant mice immunized with NMDAR1 P2 antigen. All 3 cohorts of mice were combined for analysis. **(A)** On Day 3, cued memory and fear extinction were assessed. Both groups showed robust fear extinction across trials. **(B)** Freezing percentages from CS1 were used to quantify cued memory. No differences in cued memory were detected between the two groups. **(C)** On Day 4, extinction recall was assessed. **(D)** Freezing percentages from CS1 were used to quantify extinction recall. P2-immunized mice showed significantly better extinction recall than control mice (F(1,137) = 5.09, p = 0.0256). CS0: pre-tone. Negative: Control mice; Autoantibody: IgM anti-NMDAR1 autoantibody. Data were presented as Mean+SEM. p value: * < 0.05.

## Discussion

Our studies demonstrated that blood circulating IgM anti-NMDAR1 autoantibodies against the P2 antigenic epitope are beneficial for cognitive function, in contrast to IgG anti-NMDAR1 P2 autoantibodies that impair spatial working memory in mice. Our mouse findings align with human data showing that natural plasma anti-NMDAR1 autoantibodies slow cognitive decline in patients with Alzheimer’s disease (Zhou, 2026), protect against TBI-associated depression and PTSD symptoms (Vaughn *et al*., 2026), and are associated with reduced severity of negative symptoms and improved social functioning in individuals with schizophrenia (Hansen *et al*., 2024; Luykx *et al*., 2024).

IgM anti-NMDAR1 autoantibodies are successfully induced by immunization with either complete or incomplete Freund’s adjuvant emulsified with the NMDAR1 P2 antigen. However, not all mice responded robustly to active immunization. This variability may reflect differences in the preexisting IgM autoantibody repertoire among individual Aicda mice. Further studies are needed to determine whether non- or low-responders can generate IgM anti-NMDAR1 autoantibodies effectively following booster immunization and/or at different ages. IgM anti-NMDAR1 P2 autoantibodies significantly enhance sensorimotor gating in mice, but their suppressive effect on startle amplitude varies across cohorts. It is possible that this suppressive effect is not detectable when baseline startle responses are low, but becomes evident when startle amplitudes are elevated as observed in cohort 3. Enhanced sensorimotor gating indicates more efficient sensory filtering, which supports higher-order cognitive functions. In clinical and preclinical research, better sensorimotor gating is associated with improved cognitive performance and reduced vulnerability to cognitive disruption (Bitsios *et al*, 2006; Freudenberg *et al*, 2022; Oliveras *et al*, 2015). Pro-cognitive effects of IgM anti-NMDAR1 autoantibodies were demonstrated in fear extinction recall where mice with high autoantibody levels exhibited significantly better extinction recall memory than mice with baseline levels.

About 0.1% of blood antibodies nonspecifically enter into the brain in healthy rodents and humans regardless of antibody specificities or IgG or IgM isotypes (Banks, 2010; Banks *et al*, 2007; Wang *et al*, 2018). This low-level entry likely occurs through BBB-bypassing extracellular routes and/or adsorptive-mediated transcytosis across the blood-brain barriers (Banks, 2016; Broadwell & Sofroniew, 1993). Whether such low-level penetration is sufficient to influence brain function has been debated. In our previous studies of immunized wildtype C57BL/6 mice, we found that blood IgG anti-NMDAR1 autoantibodies significantly impair mouse spatial working memory with intact blood brain barriers (Yue *et al*., 2021). Our current studies indicate that blood IgM anti-NMDAR1 autoantibodies significantly improve sensorimotor gating and fear extinction recall in mice with intact blood brain barriers. Together, these studies suggest that chronic infiltration of blood IgG or IgM anti-NMDAR1 autoantibodies may be sufficient to impact brain function. Consistent with our findings, injection of IgM anti-amyloid beta antibodies (Banks *et al*., 2007) or human natural IgM (Xu *et al*, 2015) that binds to gangliosides into mouse blood reversed mouse phenotypes of Alzheimer’s disease and amyotrophic lateral sclerosis, supporting that blood IgM antibodies can enter mouse brain to exert their functions. Such low-level brain entry may contribute to the small effect sizes observed for the pro-cognitive function of IgM anti-NMDAR1 autoantibodies on sensorimotor gating and fear extinction recall in our study. In diseases or conditions where the blood–brain barrier is compromised, autoantibodies may more readily enter the brain parenchyma, and the neuroprotective effects of IgM/natural anti-NMDAR1 autoantibodies may become more readily detectable (Hansen *et al*., 2024; Luykx *et al*., 2024; Vaughn *et al*., 2026; Zhou, 2026).

Our previous studies found that IgG anti-NMDAR1 P2 autoantibodies impairs mouse spatial working memory (Yue *et al*., 2021), but our current study suggests that IgM anti-NMDAR1 P2 autoantibodies are pro-cognitive. It is not clear how IgG and IgM anti-NMDAR1 autoantibodies against the same P2 antigenic epitope produce opposite functional outcomes in mice of the same C57 BL/6 genetic background. We hypothesize that the opposite effects of IgG vs IgM anti-NMDAR1 autoantibodies on cognitive function may be through differing effects on glutamate excitotoxicity (Zhou, 2023). Namely, IgM antibodies are physically too large to enter the synaptic cleft but selectively suppress extrasynaptic NMDAR-mediated excitotoxicity, thereby contributing to their pro-cognitive effects potentially through neuroprotection. In contrast, IgG antibodies are small enough to access the synaptic cleft where they can inhibit synaptic NMDAR function and impair neurotransmission essential for cognitive function. In the future, more experiments are needed to confirm their opposing effects, particularly electrophysiological recordings to determine whether IgM selectively suppresses extrasynaptic NMDAR currents without affecting synaptic NMDAR activity, whereas IgG suppresses both.

## Supplemental Figure Legend

**Supplemental Figure 1.**
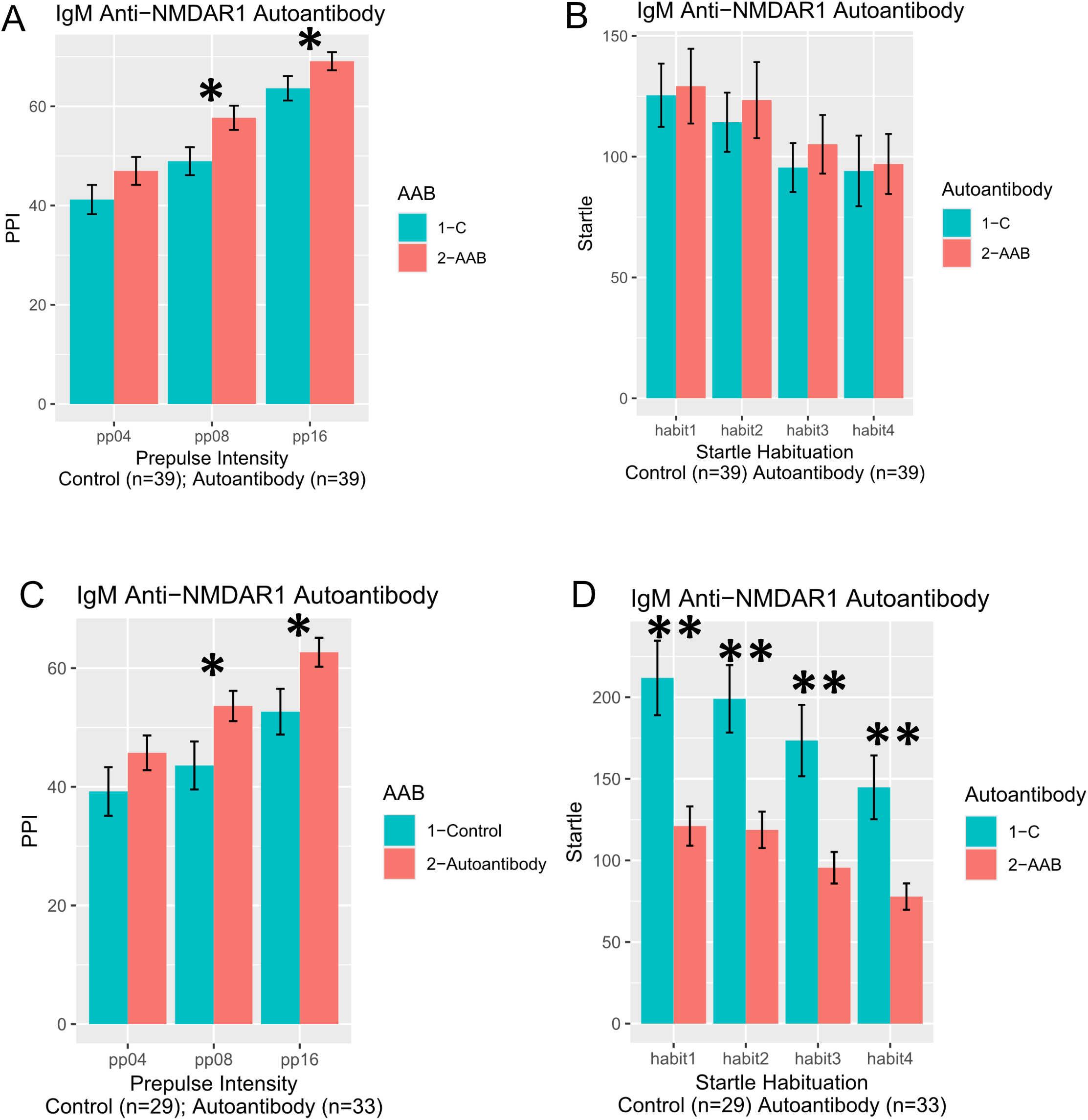
Prepulse inhibition and startle responses. Mouse cohorts 1 and 2 were combined for analysis of prepulse inhibition **(A)** and startle responses **(B)**. A significant increase in PPI was observed in mice immunized with the P2 antigen compared with control mice (F(1,74) = 4.932, p = 0.029). No differences in startle or startle habituation between the two groups. **(C)** In cohort 3, PPI was significantly increased in mice immunized with the P2 antigen compared with control mice (F(1, 58) = 4.978, p = 0.0295). **(D)** Startle amplitudes were significantly reduced in mice immunized with the P2 antigen compared with control mice (F(1,58) = 12.328, p = 0.0009). No differences in startle habituation were observed between the two groups. Data were presented as Mean+SEM. p value: ** < 0.01; * < 0.05.

## References

Anagnostaras SG, Wood SC, Shuman T, Cai DJ, Leduc AD, Zurn KR, Zurn JB, Sage JR, Herrera GM (2010) Automated assessment of pavlovian conditioned freezing and shock reactivity in mice using the video freeze system. Frontiers in behavioral neuroscience 4

Banks WA (2010) Immunotherapy and neuroimmunology in Alzheimer’s disease: a perspective from the blood-brain barrier. Immunotherapy 2:1–3

Banks WA (2016) From blood-brain barrier to blood-brain interface: new opportunities for CNS drug delivery. Nature reviews 15:275–292

Banks WA, Farr SA, Morley JE, Wolf KM, Geylis V, Steinitz M (2007) Anti-amyloid beta protein antibody passage across the blood-brain barrier in the SAMP8 mouse model of Alzheimer’s disease: an age-related selective uptake with reversal of learning impairment. Exp Neurol 206:248–256

Bitsios P, Giakoumaki SG, Theou K, Frangou S (2006) Increased prepulse inhibition of the acoustic startle response is associated with better strategy formation and execution times in healthy males. Neuropsychologia 44:2494–2499

Broadwell RD, Sofroniew MV (1993) Serum proteins bypass the blood-brain fluid barriers for extracellular entry to the central nervous system. Exp Neurol 120:245–263

Castillo-Gomez E, Oliveira B, Tapken D, Bertrand S, Klein-Schmidt C, Pan H, Zafeiriou P, Steiner J, Jurek B, Trippe R et al (2017) All naturally occurring autoantibodies against the NMDA receptor subunit NR1 have pathogenic potential irrespective of epitope and immunoglobulin class. Mol Psychiatry 22:1776–1784

Dalmau J (2016) NMDA receptor encephalitis and other antibody-mediated disorders of the synapse: The 2016 Cotzias Lecture. Neurology 87:2471–2482

During MJ, Symes CW, Lawlor PA, Lin J, Dunning J, Fitzsimons HL, Poulsen D, Leone P, Xu R, Dicker BL et al (2000) An oral vaccine against NMDAR1 with efficacy in experimental stroke and epilepsy. Science 287:1453–1460

Elkon K, Casali P (2008) Nature and functions of autoantibodies. Nat Clin Pract Rheumatol 4:491–498

Freudenberg F, Althen H, Falk K, Bittner RA, Reif A, Plichta MM (2022) Test-retest reliability of prepulse inhibition (PPI) and PPI correlation with working memory. Acta Neuropsychiatr 34:344–353

Gresack JE, Risbrough VB, Scott CN, Coste S, Stenzel-Poore M, Geyer MA, Powell SB (2010) Isolation rearing-induced deficits in contextual fear learning do not require CRF(2) receptors. Behav Brain Res 209:80–84

Hammer C, Stepniak B, Schneider A, Papiol S, Tantra M, Begemann M, Siren AL, Pardo LA, Sperling S, Mohd Jofrry S et al (2014) Neuropsychiatric disease relevance of circulating anti-NMDA receptor autoantibodies depends on blood-brain barrier integrity. Mol Psychiatry 19:1143–1149

Hansen N, Luedecke D, Maier HB, Neyazi A, Fitzner D, Wiltfang J, Malchow B (2024) NMDAR1 autoantibodies as potential biomarkers for schizophrenia phenotyping. Lancet Psychiatry 11:780–781

Jezequel J, Rogemond V, Pollak T, Lepleux M, Jacobson L, Grea H, Iyegbe C, Kahn R, McGuire P, Vincent A et al (2017) Cell- and Single Molecule-Based Methods to Detect Anti-N-Methyl-D-Aspartate Receptor Autoantibodies in Patients With First-Episode Psychosis From the OPTiMiSE Project. Biol Psychiatry 82:766–772

Ji B, Wang X, Pinto-Duarte A, Kim M, Caldwell S, Young JW, Behrens MM, Sejnowski TJ, Geyer MA, Zhou X (2013) Prolonged Ketamine Effects in Sp4 Hypomorphic Mice: Mimicking Phenotypes of Schizophrenia. PLoS ONE 8:e66327

Karpova A, Mikhaylova M, Bera S, Bar J, Reddy PP, Behnisch T, Rankovic V, Spilker C, Bethge P, Sahin J et al (2013) Encoding and transducing the synaptic or extrasynaptic origin of NMDA receptor signals to the nucleus. Cell 152:1119–1133

Lobo PI (2016) Role of Natural Autoantibodies and Natural IgM Anti-Leucocyte Autoantibodies in Health and Disease. Front Immunol 7:198

Luykx JJ, Visscher R, Winter-van Rossum I, Waters P, de Witte LD, Fleischhacker WW, Lin BD, de Boer N, van der Horst M, Yeeles K et al (2024) Clinical symptoms and psychosocial functioning in patients with schizophrenia spectrum disorders testing seropositive for anti-NMDAR antibodies: a case-control comparison with patients testing negative. Lancet Psychiatry 11:828–838

Nilaratanakul V, Chen J, Tran O, Baxter VK, Troisi EM, Yeh JX, Griffin DE (2018) Germ Line IgM Is Sufficient, but Not Required, for Antibody-Mediated Alphavirus Clearance from the Central Nervous System. J Virol 92

Oliveras I, Rio-Alamos C, Canete T, Blazquez G, Martinez-Membrives E, Giorgi O, Corda MG, Tobena A, Fernandez-Teruel A (2015) Prepulse inhibition predicts spatial working memory performance in the inbred Roman high- and low-avoidance rats and in genetically heterogeneous NIH-HS rats: relevance for studying pre-attentive and cognitive anomalies in schizophrenia. Frontiers in behavioral neuroscience 9:213

Pan H, Oliveira B, Saher G, Dere E, Tapken D, Mitjans M, Seidel J, Wesolowski J, Wakhloo D, Klein-Schmidt C et al (2019) Uncoupling the widespread occurrence of anti-NMDAR1 autoantibodies from neuropsychiatric disease in a novel autoimmune model. Mol Psychiatry 24:1489–1501

Sweeney MD, Sagare AP, Zlokovic BV (2018) Blood-brain barrier breakdown in Alzheimer disease and other neurodegenerative disorders. Nat Rev Neurol 14:133–150

Toth M, Gresack JE, Bangasser DA, Plona Z, Valentino RJ, Flandreau EI, Mansuy IM, Merlo-Pich E, Geyer MA, Risbrough VB (2014) Forebrain-specific CRF overproduction during development is sufficient to induce enduring anxiety and startle abnormalities in adult mice. Neuropsychopharmacology 39:1409–1419

Vaughn M, Acheson D, Powell S, Yurgil K, Nievergelt C, Baker D, Risbrough V, Zhou X (2026) Potential Neuroprotective Effects of Natural Anti-NMDAR1 Autoantibodies Against Psychiatric Symptoms Associated with Traumatic Brain Injuries. medRxiv: 2026.2001.2008.26343690

Vaughn M, Powell S, Risbrough V, Zhou X (2025) A novel simple immunoassay for quantification of blood anti-NMDAR1 autoantibodies. PeerJ 13:e19212

Wang Q, Delva L, Weinreb PH, Pepinsky RB, Graham D, Veizaj E, Cheung AE, Chen W, Nestorov I, Rohde E et al (2018) Monoclonal antibody exposure in rat and cynomolgus monkey cerebrospinal fluid following systemic administration. Fluids Barriers CNS 15:10

Wu QJ, Tymianski M (2018) Targeting NMDA receptors in stroke: new hope in neuroprotection. Molecular brain 11:15

Xia P, Chen HS, Zhang D, Lipton SA (2010) Memantine preferentially blocks extrasynaptic over synaptic NMDA receptor currents in hippocampal autapses. J Neurosci 30:11246–11250

Xu X, Denic A, Jordan LR, Wittenberg NJ, Warrington AE, Wootla B, Papke LM, Zoecklein LJ, Yoo D, Shaver J et al (2015) A natural human IgM that binds to gangliosides is therapeutic in murine models of amyotrophic lateral sclerosis. Dis Model Mech 8:831–842

Yue W, Caldwell S, Risbrough V, Powell S, Zhou X (2021) Chronic presence of blood circulating anti-NMDAR1 autoantibodies impairs cognitive function in mice. PLoS ONE 16:e0256972

Zhou X (2021) Cognitive Impact by Blood Circulating Anti-NMDAR1 Autoantibodies. J Psychiatr Brain Sci 6

Zhou X (2023) Preventive and Therapeutic Autoantibodies Protect against Neuronal Excitotoxicity. J Psychiatr Brain Sci 8

Zhou X (2026) Natural Anti-NMDAR1 autoantibodies associate with slowed decline of cognitive functions in Alzheimer’s diseases. Translational psychiatry 16

